# Effects of arbuscular mycorrhizal fungi on nitrogen uptake in cotton (*Gossypium hirsutum* L.) under low-nitrogen conditions

**DOI:** 10.1101/2024.03.17.585399

**Authors:** Hushan Wang, Yijian Wang, Xiaojiao Cheng, Yunzhu He, Zihui Shen, Wangfeng Zhang, Xiaozhen Pu

## Abstract

**Summary:**

- Cotton is an important global cash crop whose yield and quality are highly influenced by soil nitrogen. Therefore, examining the interactions between roots and arbuscular mycorrhizal fungi (AMF) under reduced nitrogen conditions is of great significance.
- We investigated the effects of nitrogen application (0, 250, and 375 kg· hm^-2^) on the AMF infection rate of cotton, the nitrogen content of each organ, root morphological characteristics and biomass, soil extracellular enzyme activity, and soil carbon and nitrogen content using a compartmentalized culture system.
- The contribution of AMF to plant nitrogen was 10.40, 22.72, and 16.67% under high, low, and no nitrogen treatments, respectively. Under low-nitrogen conditions, the symbiosis between AMF and roots increased root surface area, tip number, branch number, mean diameter, and biomass; and increased soil extracellular enzyme activity (protease, NAG, PER, and PPO), the microbial biomass carbon-to-nitrogen ratio, active carbon content, and the soil nitrogen mineralization rate. Soil NO_3_^-^-N, NH_4_^+^-N, and organic nitrogen content decreased, whereas the absorption of NO_3_^-^-N by AMF hyphae was higher than that of NH_4_^+^-N.
- Under low-nitrogen conditions, AMF promoted the decomposition of soil organic matter and the transformation of soil nitrogen through the action of hyphal microorganisms.

## Introduction

Cotton (*Gossypium hirsutum* L.) is a fiber crop with the largest cultivated area and widest distribution worldwide, with China being the largest producer and consumer of cotton. Nitrogen is the most important mineral element in plants and determines crop yield and quality (Zhou *et al*., 2021). This greatly limits crop productivity owing to the widespread nitrogen deficiency in Chinese farmland soils. In China, high cotton yields have been achieved through intensive water and nitrogen fertilizer use; however, only 25–50% of the nitrogen fertilizer applied to farmland soil is absorbed and utilized by crops, whereas much of the rest is lost from the soil/plant system. This reduces soil fertility and crop yields and leads to a series of ecological problems (Zaman & Blennerhassett, 2010; Wang *et al*., 2021). Therefore, a thorough understanding of the characteristics and mechanisms of nitrogen uptake and utilization by plants and the improvement of soil nitrogen supply capacity are prerequisites for efficient nutrient utilization and regulation in farmland ecosystems in China.

Arbuscular mycorrhizal fungi (AMF) can establish mutualistic symbionts with more than 80% terrestrial plants, including most crops (Brundret *et al*., 1996). As the primary way of plant nutrient acquisition, many studies have shown that the symbiosis between root systems and AMF can activate soil nutrients and promote nitrogen absorption (Halder *et al*., 2015). This is because AMF epitaxial hyphae can improve the soil microenvironment by secreting a series of carbon-containing compounds or releasing photosynthetic carbon into the soil through the turnover of the hyphae and also serve as a food source for soil microorganisms that stimulate nitrogen transformation. Ultimately, AMF act as a nutrient activator. Simultaneously, AMF can directly affect plant nitrogen uptake (via the hyphae pathway) through extracellular hyphae uptake and intra-root hyphae transport processes (Smith & Smith, 2011), as well as indirectly affect nitrogen uptake by changing root morphology and physiology and regulating the expression of nitrogen-related transporters (Hobbie & Hobbie, 2008). In addition, AMF are able to fix a large amount of nitrogen in their mycelia and can act as a nitrogen sink, thereby reducing nitrogen leaching (Gilliam, 2006). Moreover, many studies have suggested that AMF infection can positively impact crop growth and biomass by promoting nitrogen absorption (Hawkins *et al*., 2000; Stavros *et al*., 2011; Waller *et al*., 2018). Therefore, AMF have the potential to activate soil nutrients, improve plant nutrition, and reduce nutrient loss, particularly in nutrient-deficient soils.

AMF can have a positive effect on nitrogen absorption by plants. The reasons for this are as follows. First, AMF symbiosis promotes plant carbon assimilation enzyme activity, increases the leaf carbon source and photosynthetic pigment content, improves light energy use efficiency, and increases underground carbohydrate input. This results in the continuous extension of the hyphal network and root system, thereby, the nutrient absorption area is expanded (Yang *et al*., 2016; Jiang *et al*., 2017). Second, AMF regulate nitrogen uptake by roots by directly affecting root growth Jabborova *et al*., 2021). Third, the life-cycle activities of AMF hyphae (e.g., hyphal secretions and dead hyphal tissues) indirectly affect the decomposition of organic carbon and the transformation of nitrogen by influencing the growth and activity of hyphal microorganisms (Dijkstra *et al*., 2021). Finally, AMF can promote nitrogen transfer between adjacent plants through underground hyphal networks (“hyphal bridges”) and adjust the soil nitrogen distribution according to plant nitrogen requirements (Van der Heijden, 2001). Therefore, the symbiosis between AMF and plants significantly and profoundly affects N absorption by plants. Nevertheless, the effect of AMF on nitrogen is influenced by many factors, which greatly increase the uncertainty regarding the effect of AMF on plant nitrogen (Guo *et al*., 2018). For example, recent studies have shown that, owing to differences in AMF composition, quantity, and basic soil characteristics, AMF have different absorption preferences for different forms of soil nitrogen (NO_3_^-^-N and NH_4_^+^-N) (Pan *et al*., 2020). It is generally believed that AMF prefer obtaining NH_4_^+^-N from the soil (Hodge, 2001). Seck-Mbengue *et al*. (2017) showed that AMF absorbed and transported twice as much nitrogen to plants under ammonium nitrogen fertilizer treatment as under nitrate nitrogen fertilizer. However, Wang *et al*. (2020) found that the symbiotic transport of NH_4_^+^-N may not be the only condition for the symbiotic transfer of nitrogen nutrients in AMF, and that the NO_3_^-^-N absorption pathway exists widely in AMF symbiosis, which may be related to plant species, AMF community composition, and soil nitrogen application level (form and content). Wang *et al*. (2023) showed that AMF inoculation is conducive to NO_3_^-^-N absorption when nitrogen is high but is conducive to NH_4_^+^-N absorption when nitrogen is low.

The C and N cycles in terrestrial ecosystems show strong coupling relationships, including C assimilation during plant photosynthesis, the decomposition of soil organic matter, root turnover, respiration, and other C-release processes. Nitrogen mineralization is an important part of the plant–soil C–N nutrient cycle and a key factor in regulating mycorrhizal symbiont interactions. Hodge and Fitter (2010) found that AMF enhance the degradation of organic matter mainly by recruiting more soil free microorganisms to the mycelium, including bacteria related to nitrogen mineralization. Driven by these microorganisms, soil enzyme-mediated catalysis improves the mineralization rate of organic matter. Thus, nitrogen storage in the soil can be mined under nitrogen-limited conditions to promote plant growth and development (Pellitier & Zak, 2018; Fu *et al*., 2022). At the same time, in return, plants also provide carbon sources to AMF (Smith & Read, 2008), and this process is affected by biological and abiotic factors such as soil physicochemical properties, microorganisms, and root growth and development (Bowles *et al*., 2015). In addition, AMF can promote the release of N and the accumulation of active C in soil organic matter—through the inter-hyphal excitation effect—to meet the nutritional demands of plants and microorganisms (Zhang *et al*., 2014; See *et al*., 2022). Xinjiang has become the main cotton-producing area in China owing to its climatic advantages; however, low nitrogen use efficiency is a key limiting factor in the further improvement of cotton yield and quality in Xinjiang. The use of AMF has been proposed as an effective solution to this problem. Here, we hypothesized that AMF and root symbiosis could improve the nitrogen-absorption capacity of cotton and that this effect is influenced by different nitrogen application amounts. Based on our experiments, we show that the lower the nitrogen level, the greater the promoting effect of AMF. Thus, our results provide a theoretical basis for improving the quality, efficiency, and yield of cotton production in other cotton-producing areas.

## Materials and Methods

### Test materials

The soil used in our experiment was obtained from the Second Company (85°59 ‘42’E, 44°19‘19’N) of the Shihezi University Test Site, Xinjiang, China. The average annual precipitation in this region is 210.6 mm; the average evaporation is 1,664.1 mm; the frost-free period is approximately 170 d; the average annual temperature is 7.0 ℃; and the annual sunshine duration is 2,861.2 h. The tested cotton variety was Xinluzao 84. The experimental soil type was irrigated, gray desert soil with a loam texture (Table 1).

**Table 1.** Physical and chemical properties of the experimental soil.

| pH | Organic<br>matter<br>(g·kg<sup>-1</sup>) | Total<br>nitrogen<br>(g·kg<sup>-1</sup>) | Available<br>phosphorus<br>(mg·kg<sup>-1</sup>) | Available<br>potassium<br>(mg·kg<sup>-1</sup>) | Alkali hydrolyzed<br>nitrogen (mg·kg<sup>-1</sup>) |
| --- | --- | --- | --- | --- | --- |
| 8.31 | 7.70 | 0.97 | 20.10 | 250.65 | 40.98 |

### Experimental design

We used a two-factor experimental design. Factor 1 was nitrogen application treatment with three levels, i.e., N1 (the typical application rate used for the drip irrigation of cotton under film in Xinjiang fields, 375 kg·hm^-2^); N2 (2/3 of N1, 250 kg·hm^-2^); and N3 (no nitrogen application). Nitrogen fertilizer was applied with water using a syringe. Factor 2 was AMF treatment, which aimed to distinguish between the direct effects of AMF and the joint effects of root mycelia. For this, 1.2-mm-thick iron plates were used to fabricate a device (13 × 10 × 13 cm) with the middle portion separated using nylon mesh; one side formed the cotton growth chamber and the other side the mycelium chamber. Nylon mesh with two different pore sizes (0.45 and 38 μm, respectively) was used across three levels, i.e., F_1_^+^ (soil containing AMF, nylon mesh aperture = 38 μm, which allowed AMF hyphae to enter the hyphae chamber but not the root system); F_0_^+^ (soil containing AMF, nylon mesh aperture = 0.45 μm, through which neither the root system nor the AMF hyphae would enter the hyphae chamber, growing only in the growth chamber); and F_0_^-^ (soil without AMF, nylon mesh aperture = 0.45 μm) (Fig. 1). This design gave nine levels of treatment in total, with each process repeated four times.

**Fig. 1.**
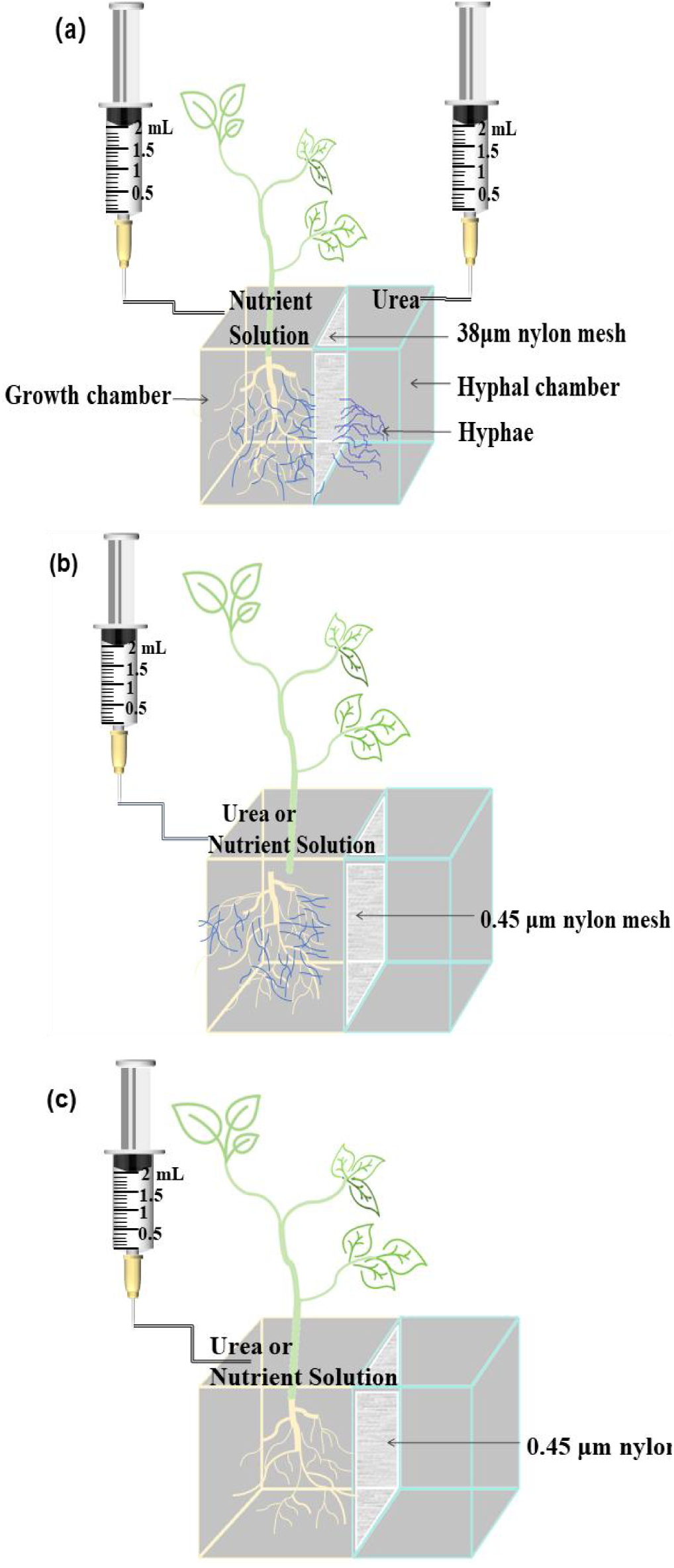
Experimental design of the AMF treatment: (a) F_1_^+^, (b) F_0_^+^, (c) F_0_^-^.

Selected plump cotton seeds were soaked in 5% H_2_O_2_ for 15 min and rinsed with running water. The growth and hyphal chambers were loaded with 600 and 1,200 g of soil, respectively. The F_0_^+^ and F_1_^+^ treatments included fresh soil (with AMF). In the F_0_^-^ treatment, fresh soil was autoclaved and loaded with 10 mL of soil filtrate without AMF to restore the soil microflora before sterilization. Four seeds were planted per growth chamber. Subsequently, the whole device was cultured in a GXZ-430D intelligent light incubator with a daily light intensity of 24,000 lx for 16 h at 28 °C and darkness for 8 h at 25 °C. An appropriate amount of deionized water was added daily. After two true cotton leaves had developed, two plants with the same growth potential and uniform spacing were maintained in the growth chamber. Two milliliters of the corresponding nitrogen level was then applied to the hyphae chamber for treatments N1 and N2 every 2 d. For N3, the same volume of deionized water was applied in place of nitrogen. Nitrogen was added as urea. Simultaneously, 2 mL of modified Hoagland non-nitrogen nutrient solution was added to the growth chambers (Fig. 1). After 60 d, the plants were harvested and observed.

## Sample collection and index determination

### Sample collection

After measuring the height of the seedlings using a tape measure, the roots, stems, and leaves were harvested. A portion of the root samples was stored in the refrigerator at 4 ℃ for the determination of the mycorrhizal infection rate and nutrient and morphological indices. The roots used to determine mycorrhizal infection rates were fixed in a standard fixing solution. The remaining roots, stems, and leaves were put in envelopes and placed in an oven at 105 ℃ for 30 min and then placed in an oven at 70 ℃ for 48 h for the determination of plant nitrogen content and biomass. Simultaneously, soil samples from each treatment were collected, passed through a 2-mm screen, and stored in plastic zip-lock bags in a refrigerator at 4 ℃ for the determination of soil extracellular enzyme activity and nutrient content.

### Index measurement

Using an EPSON EXPRESSION 10000XL root scanner (Seiko Epson Corp., Japan) and WinRHI-ZO software (Pro 2013e, Regent Instruments Inc., Canada), total root length, surface area, mean diameter, volume, number of tips, and number of branches were determined. NO_3_^-^-N, NH_4_^+^-N, and total nitrogen in the plant roots were determined using salicylic acid–sulfuric acid colorimetry (Li & Zhang, 2016), ultraviolet spectrophotometry (Sun *et al*., 2013), and Nessy colorimetry (Bao, 2000).

Soil microbial biomass carbon (MBC) and microbial biomass nitrogen (MBN) were determined using the chloroform fumigation method (Lu, 2000). Soil chitinolytic enzyme (β-N-acetyl-glucosaminidase, NAG), lignin-degrading enzyme (peroxidase, PER, and polyphenol oxidase, PPO), and protease were evaluated using ultraviolet spectrophotometry (Parham & Deng, 2000), iodine liquid titration, and sodium caseinate colorimetry, respectively (Zheng *et al*., 1980). Soil NH_4_^+^-N and NO_3_^-^-N content was determined using phenol disulfonic acid and indigophenol blue colorimetry, respectively (Bray & Kurtz, 1945). The free amino acid (FAA) content of the soil was determined using ninhydrin colorimetry (Joergensen, 1996). Soil dissolved organic nitrogen (DON) content was determined using the reduction method of Dalton alloy (State Environmental Protection Administration in China, 2002). The soil N mineralization rate and the FAA–net amino acid production rate (FAA–NPR) were measured by indoor culture (Ross *et al*., 1999). The content of soil readily oxidizable carbon (ROC) and total organic carbon (TOC) were determined using 333 mmol·L^-1^ potassium permanganate oxidation and external heating of potassium dichromate, respectively (Blair *et al*., 1995). The inert organic carbon (IOC) content of the soil was subsequently determined as the difference between the TOC and ROC (Blair *et al*., 1995).

The roots were then stained with Trypan Blue. The root segments in the standard fixing solution were removed and washed three to five times with distilled water, cut into 1.0–1.5-cm segments, placed in 10% (mass fraction) KOH at 90 ℃ for 20 min, and then rinsed with distilled water three to five times. Decolorization was performed in 10% H_2_O_2_ for 3 h and then washed three to five times with distilled water. Acidizing was performed using 2% hydrochloric acid at room temperature for 20 min, followed by dyeing with 0.05% Trypan Blue in a water bath at 90 ℃ for 20 min, rinsing the samples with distilled water three to five times. Decolorization was performed in lactate glycerol solution for 1 h. The root segments were removed and treated with polyvinyl alcohol lactate glycerol (PVLG) for observation. Thirty root segments were assigned randomly to each treatment group. The cross method was used to count infected mycorrhizal root segments (Giovannetti & Mosse, 2010), and the AMF infection rate was calculated as follows:

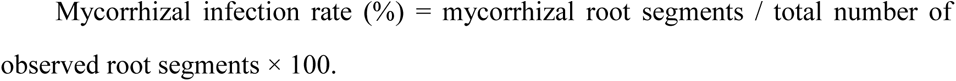

In addition, the contribution rate of AMF hyphae was calculated as follows:

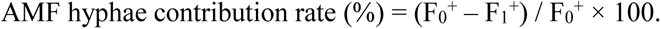

The difference of inorganic nitrogen uptake by roots was calculated as follows: Inorganic nitrogen-absorption difference of roots = root NO_3_^-^-N – root NH_4_^+^-N

### Data Analysis

Excel 2019 (Beijing Kingsoft Office Software Co., Ltd) and SPSS 23.0 (Statistical Product and Service Solutions; IBM company) were used for data processing and analysis. Before the analysis, a variance homogeneity test was performed, and logarithmic conversion was applied when homogeneity was not met. Nitrogen application, AMF, and their interactions were evaluated using a two-factor ANOVA. Duncan’s multiple comparison test was used to compare the mean values between the groups. The significance level was set at *P* = 0.05. All plots were produced using Origin 2024 (OriginLab Corporation).

## Results

### Influence of nitrogen application and AMF on cotton root infection rate

Regardless of the nitrogen application level, no AMF infection was observed in the F_0_ ^-^ treatment (Fig. 2c). AMF and cotton roots formed mycorrhizal symbiosis under the F_0_^+^ and F ^+^ treatments (Fig. 2a,b). With a decrease in the nitrogen application level, the AMF infection rate of the cotton roots gradually increased. At the N3 level, the AMF infection rate of the cotton roots was the highest, reaching 83.33%. At the N1 level, the infection rates of the F_0_^+^ and F_1_^+^ treatments were 73.33% and 66.67%, respectively, compared to 80% and 80% at the N2 level (Table 2).

**Fig. 2.**
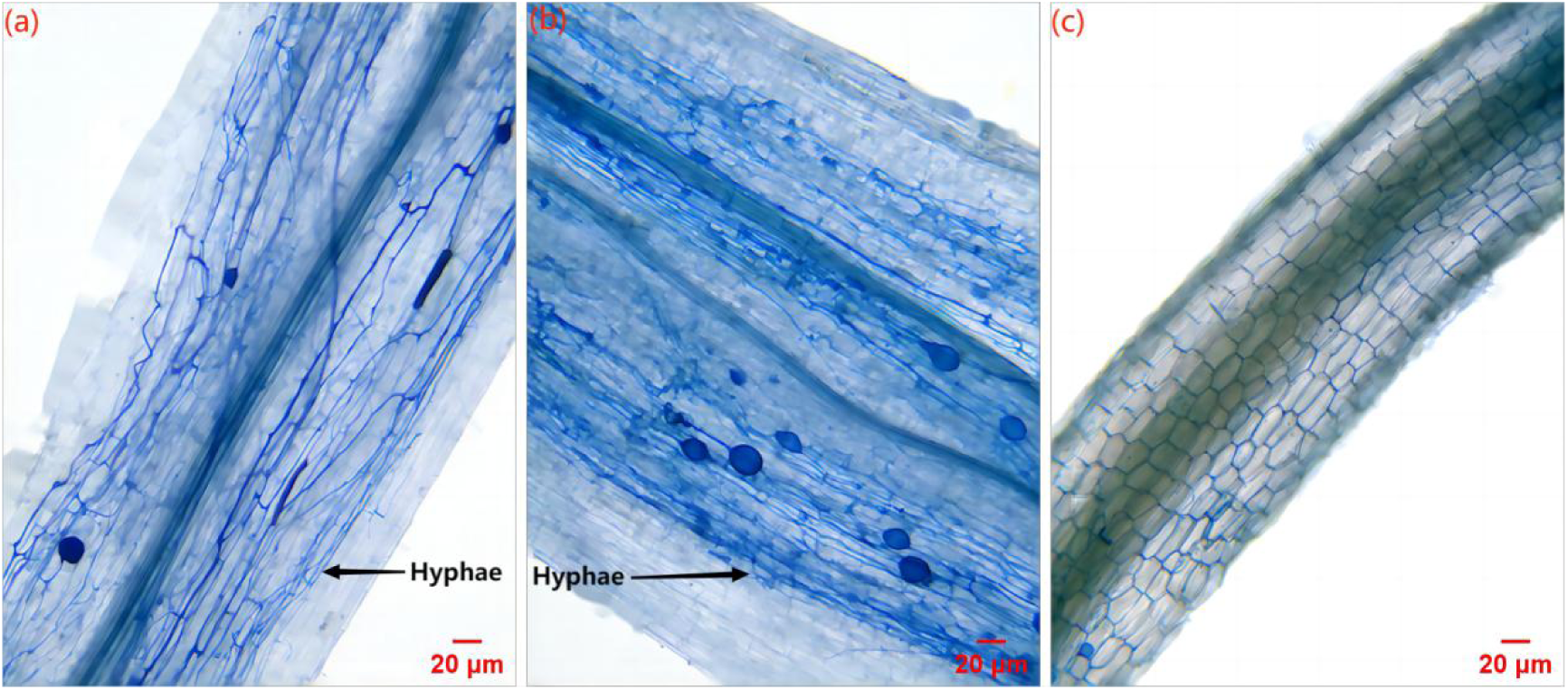
Infection of AMF on cotton roots under different nitrogen application levels: (a), (b), and (c) are F_0_^+^, F_1_^+^, and F_0_ ^-^, respectively.

**Table 2.**
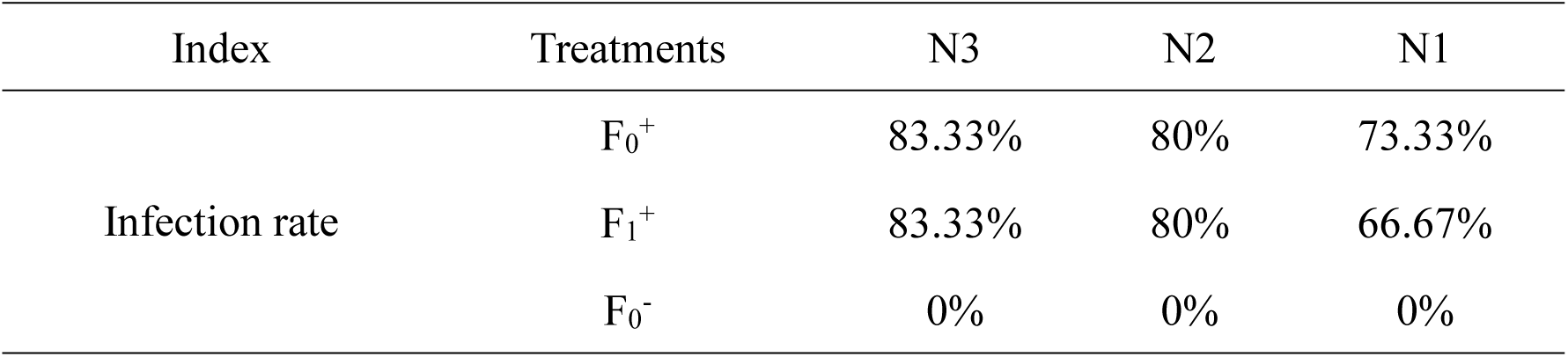
Infection rates of cotton roots by AMF under different nitrogen application levels.

### Effects of nitrogen application and AMF on cotton plant height, biomass, and root morphology

Fig. 3a shows that the height of plants treated with F_1_^+^ was significantly higher than that of plants treated with F_0_^+^ and F_0_^-^ under the same nitrogen application level. There was no significant difference between F_0_^+^ and F_0_^-^ at the N2 and N1 levels; however, there was a significant difference under conditions of N3 treatment. Under the N2 and N1 levels, the total root length of F_0_^+^ and F_1_^+^ was lower than that of F_0_^-^, but the root surface area, average diameter, volume, number of tips, and branching number were all higher than those of F_0_^-^ (Table 3). This effect was markedly more significant at the N2 level.

**Fig. 3.**
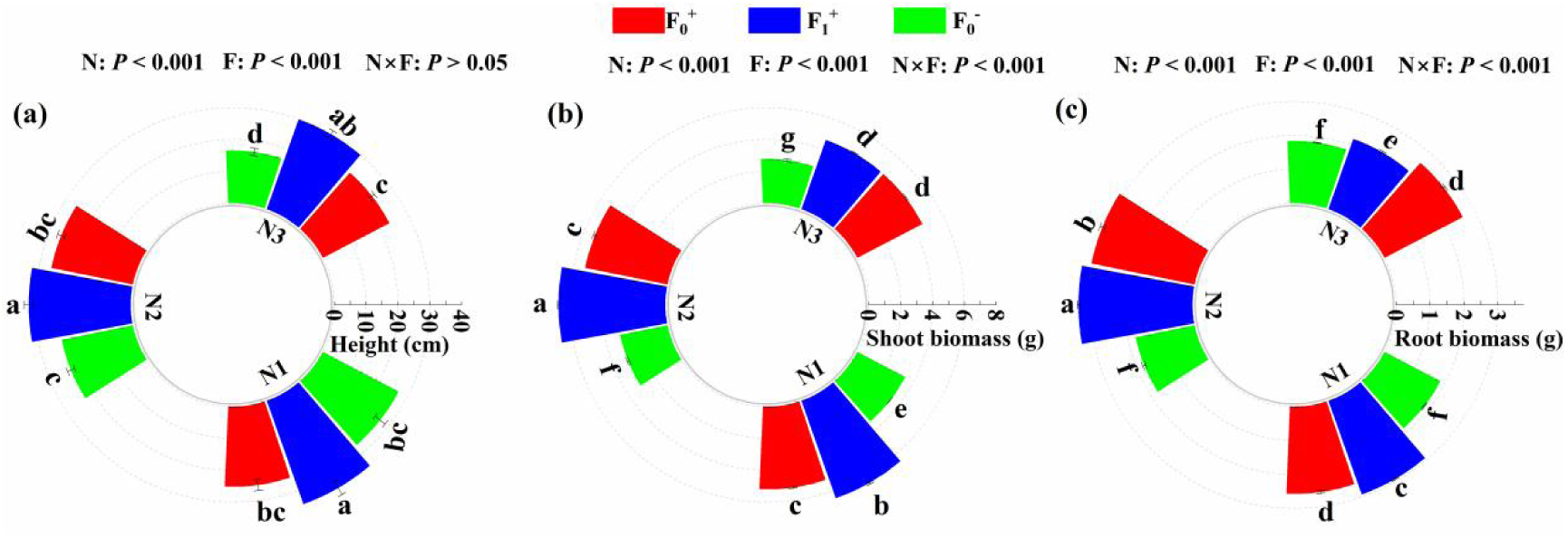
Effects of AMF on cotton plant height (a), shoot biomass (b), and root biomass (c) under different nitrogen application levels.

**Table 3.**
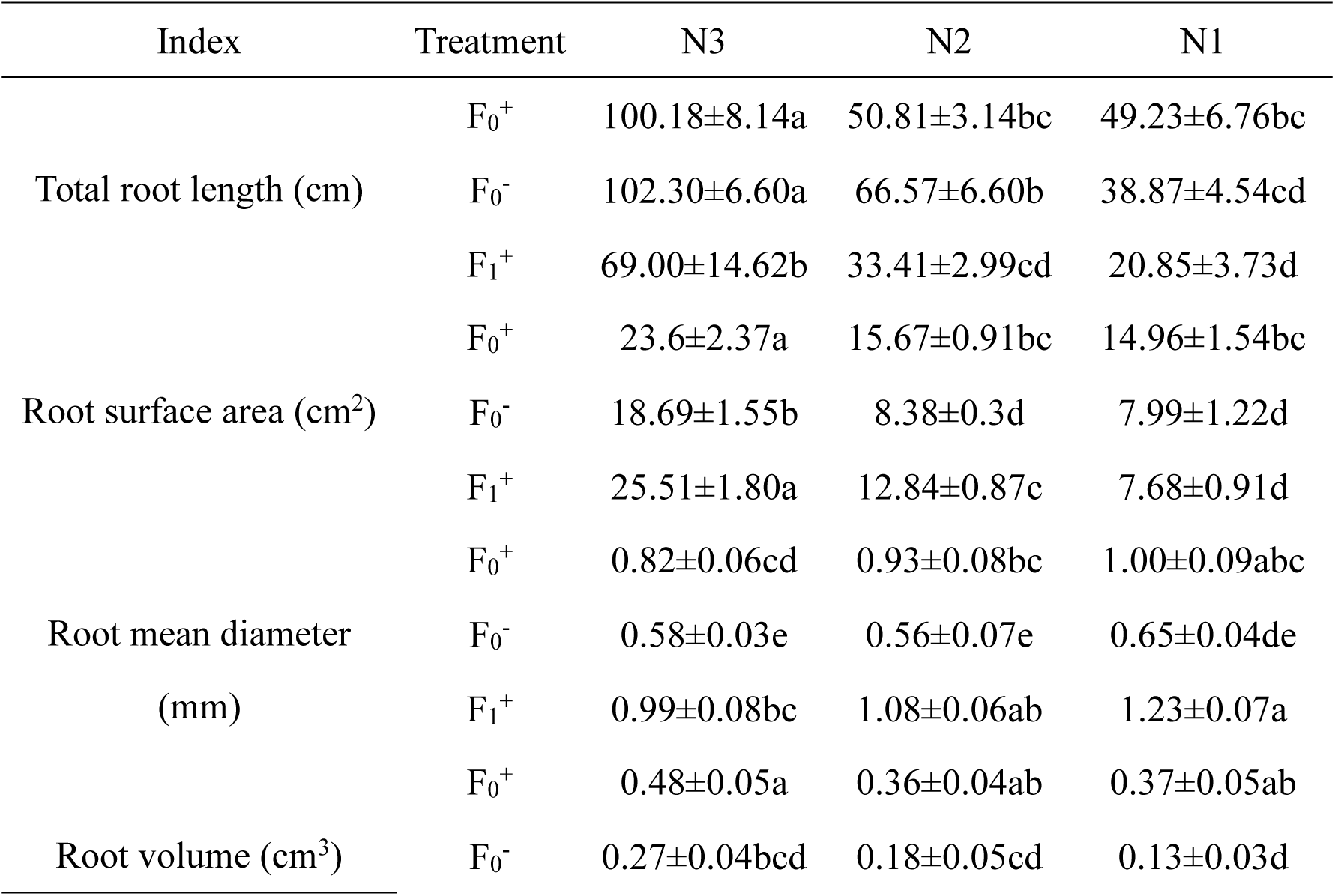
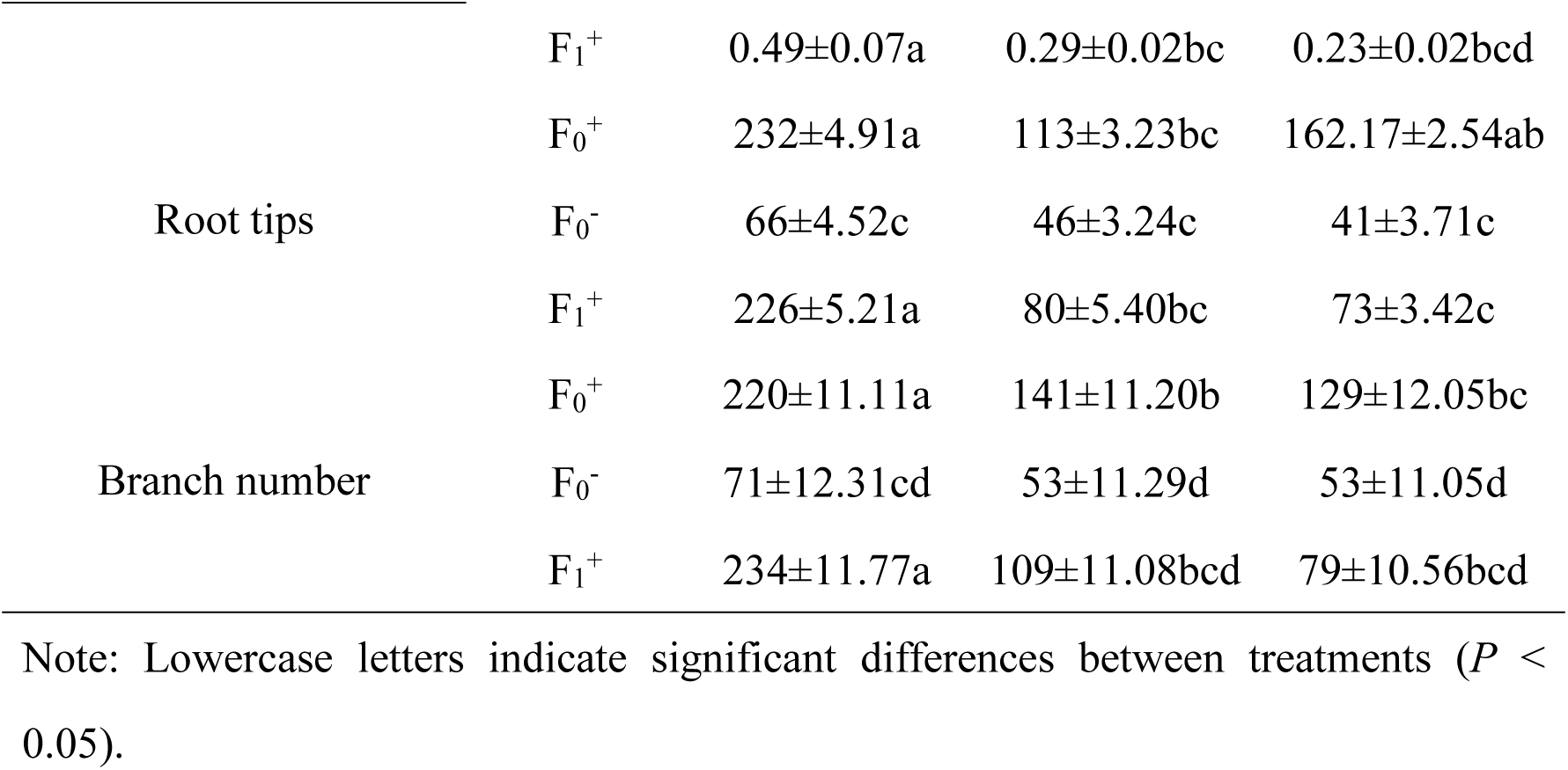
Effects of AMF on cotton root morphology under different nitrogen application levels (mean ± standard error).

Under the same nitrogen application level, F_0_^+^ and F_1_^+^ significantly increased cotton shoot and root biomass, and F_1_^+^ was significantly higher than F_0_^+^ (Fig. 3a,b). At the N1 and N2 levels, F_0_^+^ and F_1_^+^ significantly increased root and shoot biomass compared to F_0_^-^, and the increase was greater at the N2 level.

### Effects of nitrogen application and AMF on the total nitrogen content of cotton organs

With an increase in nitrogen application, the total nitrogen content of cotton roots and aboveground parts changed significantly; however, the total nitrogen content of cotton roots was higher than that of the aboveground parts, with a difference of 0.02–105.91 mg·g^-1^ (Fig. 4a,b). At the same nitrogen level, the root total nitrogen content of F_0_^+^ was higher than that of F_1_^+^, and the root total nitrogen content of F_0_^+^ was significantly higher than that of F_1_^-^. The contribution rate of AMF mycelia to the total nitrogen content of cotton differed under the different nitrogen levels. Specifically, the contribution rate was higher under conditions of N1 treatment (Fig. 4b).

**Fig. 4.**
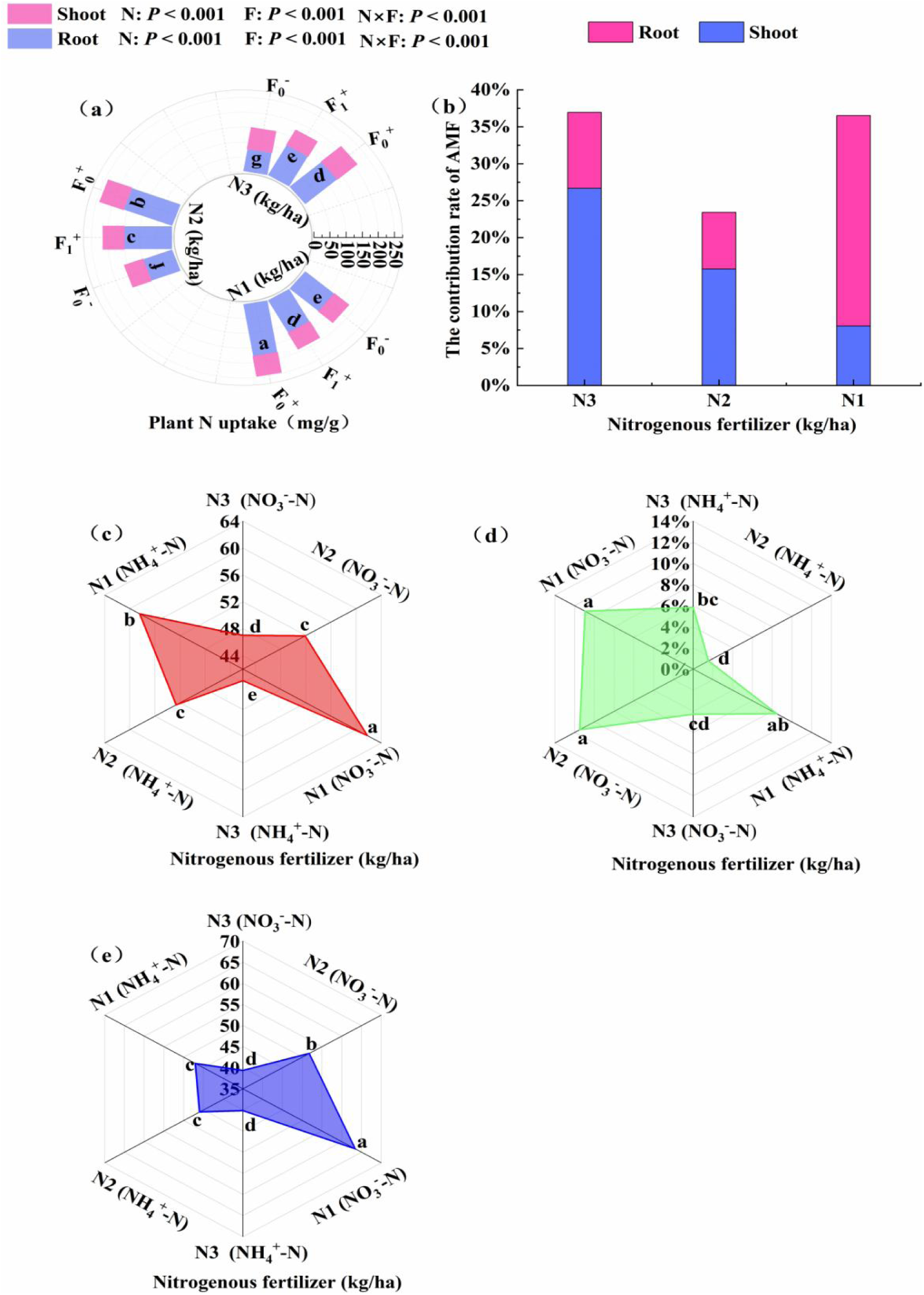
Effects of AMF on nitrogen content in cotton under different nitrogen application levels: (a) differences in total nitrogen content between aboveground and roots; (b) contribution rate of mycelium to nitrogen in shoots and roots; (c) influence of mycelium on the NO_3_^-^-N and NH_4_^+^-N content of roots; (d) the contribution rate of mycelium to the relative contents of NO_3_^-^-N and NH_4_^+^-N in roots; (e) changes in NO_3_^-^-N and NH_4_^+^-N in roots without AMF.

### Effects of nitrogen application and AMF on the inorganic nitrogen content of cotton roots

The inorganic nitrogen content of the roots was significantly higher under N1 and N2 levels than that under the N3 level (Fig. 4e). Under the same nitrogen application level, the root NO_3_^-^-N content was significantly higher than that of NH_4_^+^-N, and the nitrogen content of the roots treated under F_0_^+^ was higher than that of F_1_^+^. Under the different nitrogen application levels, the contributions of AMF mycelia to NO_3_^-^-N and NH_4_^+^-N in the cotton roots were significantly different. With an increase in nitrogen application, the contributions of both NO_3_^-^-N and NH_4_^+^-N to the root system by AMF mycelia increased, with the highest contribution rate at 11.49% and the lowest at 1.55%. The contribution of AMF mycelia to NO_3_^-^-N was greater than that to NH_4_^+^-N (Fig. 4d). In addition, with an increase in the nitrogen application rate, NO_3_^-^-N and NH_4_^+^-N in F_0_^-^-treated roots both increased. The NO_3_^-^-N content of the roots was higher than the NH_4_^+^-N content, with a difference of 0–16 mg·g^-1^ (Fig. 4e).

### Effects of nitrogen application and AMF on soil extracellular enzymes

With an increase in nitrogen application, soil extracellular enzyme activity decreased significantly (Fig. 5). Under the same nitrogen application level, the activities of soil protease, NAG, PER, and PPO in the F_1_^+^ and F ^+^ treatments were significantly increased compared with those in F_0_^-^. These changes were more pronounced in the N2 treatment (Fig. 5). At the N2 level, the activities of the four enzymes in the F_1_^+^ treatment were significantly higher than those in F ^+^. Regardless of the AMF treatment, enzyme activity at the N1 level was the lowest among the three tested levels.

**Fig. 5.**
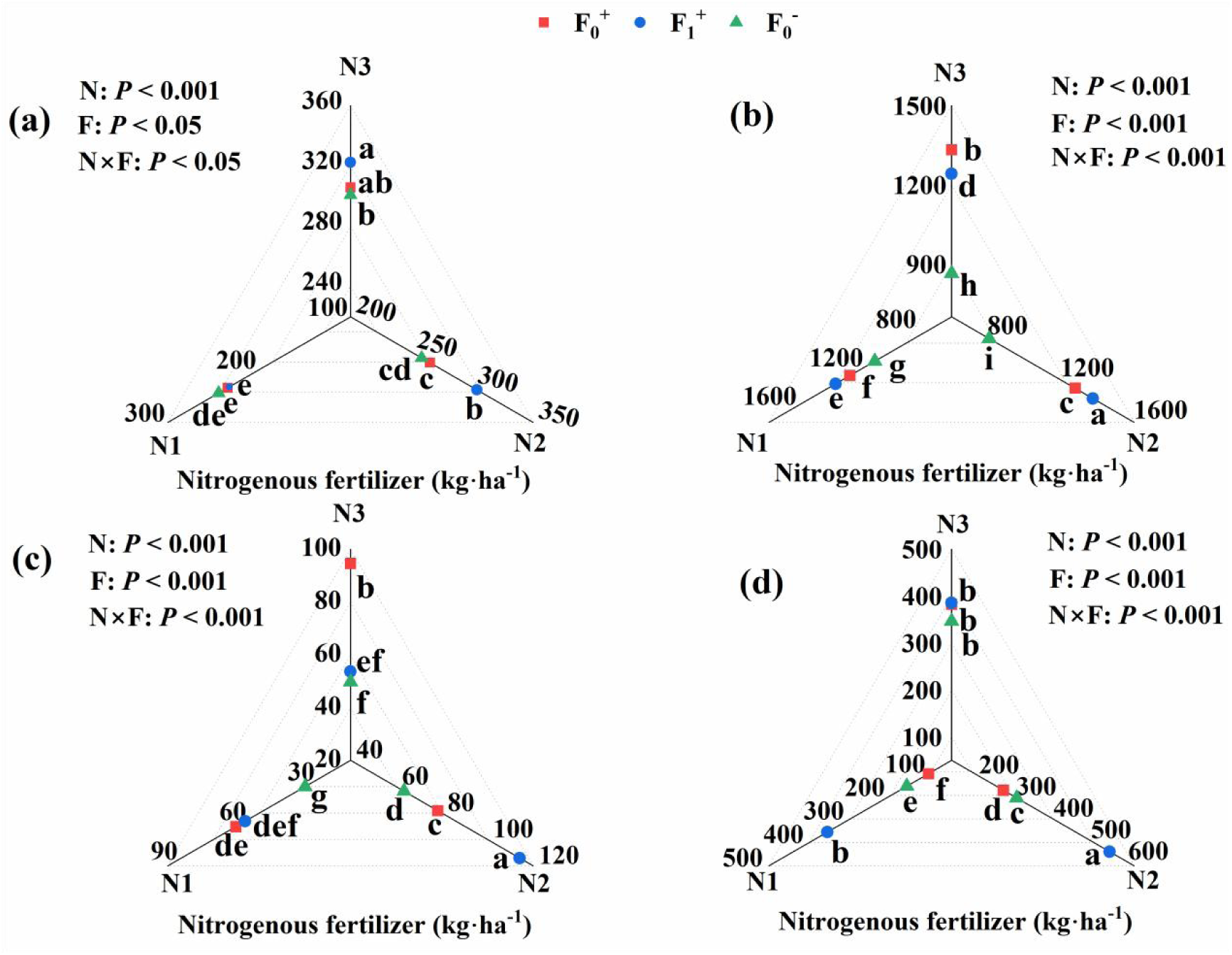
Effects of AMF on soil protease activity (a), β-N-acetylglucosamine glycosidase activity (b), peroxidase activity (c), and polyphenol oxidase activity (d) under different nitrogen application levels.

### Effects of nitrogen application and AMF on soil inorganic nitrogen, FAA, and DON

At the same nitrogen application level, the soil NO_3_^-^-N content of F_0_^+^ and F_1_^+^ decreased compared to that of F_0_^-^, and there was a significant difference between the F_0_^+^ and F_1_^+^ treatments (Fig. 6a). At the N3 level, the NO_3_^-^-N content of the F_0_^+^ treatment group was significantly higher than that of the F_1_^+^ treatment group. At the N2 level, the NO_3_^-^-N content of the soil treated with F_1_^+^ was significantly higher than that in the soil treated with F_0_^+^. At the N1 level, there was no significant difference in NO_3_^-^-N content between the F_1_^+^ and F_0_^+^ treated soils. Under the same nitrogen application level, the NH_4_^+^-N content of the soil was generally higher than the NO_3_^-^-N content (Fig. 6b), and the NH_4_^+^-N content in F_1_^+^ was significantly lower than that in F_0_^+^ and F_0_^-^. At the N2 level, there was no significant difference in the NH_4_^+^-N content between the F_0_^-^ and F_0_^+^ treatments. At the N1 level, the NH_4_^+^-N content of the F_0_^-^ treatment was significantly higher than that of F_0_^+^. Under the same nitrogen application level, the NO_3_^-^-N mineralization rate of the F_1_^+^ treatment was the highest and that of the F_0_^+^ treatment was the lowest (Fig. 6c). In addition, with an increase in nitrogen application, the mineralization rate of NH_4_^+^-N gradually decreased (Fig. 6d).

**Fig. 6.**
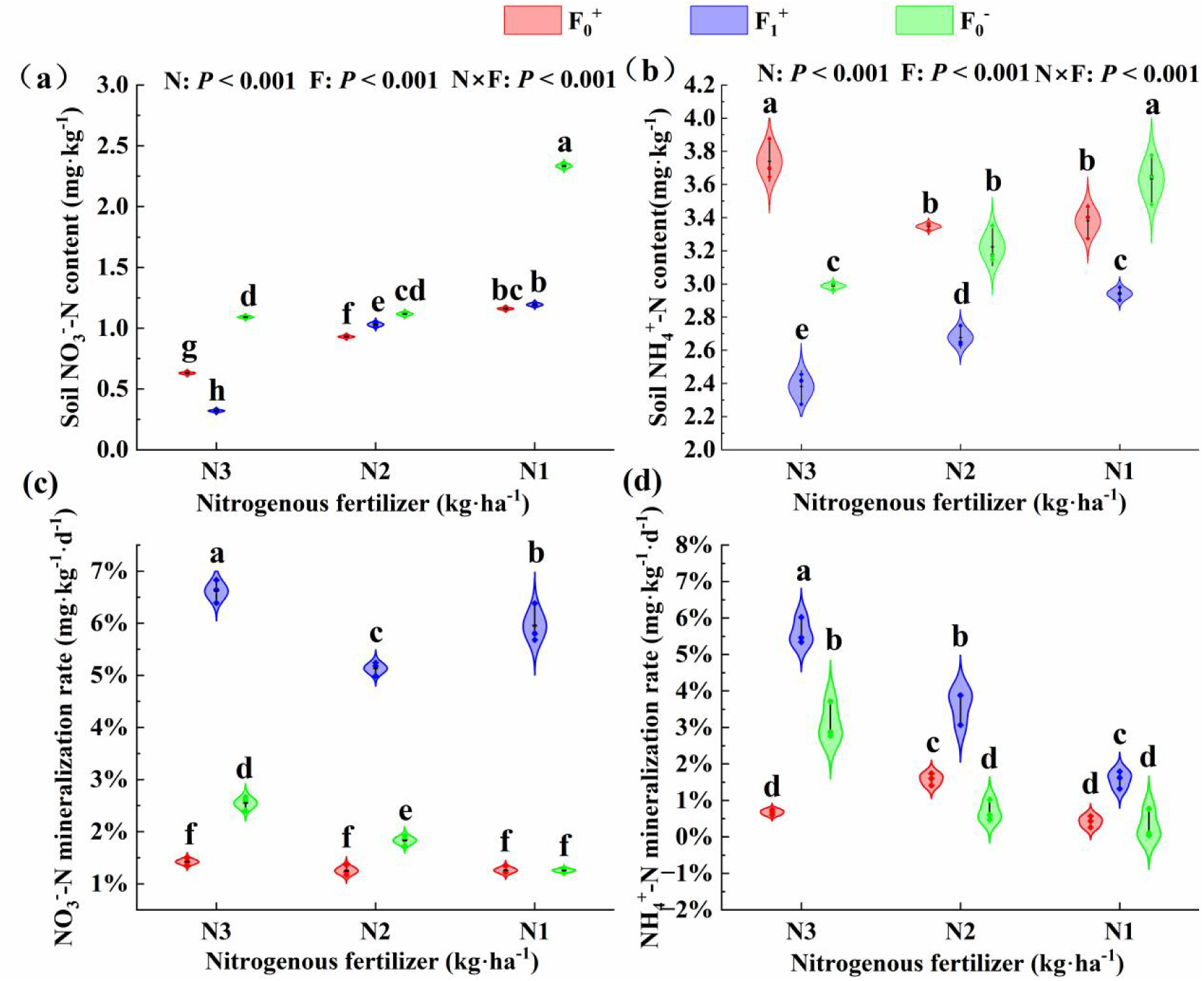
Effects of nitrogen application and AMF on (a) NO_3_^-^-N content, (b) NH_4_^+^-N content, (c) NO_3_^-^-N mineralization rate, and (d) NH_4_^+^-N mineralization rate in soil.

At the same nitrogen application level, the FAA content of the F_0_^+^ and F_1_^+^ soils was lower than that of the F_0_^-^ treatment (Fig. 7a), but the FAA–NPR of the F_0_^+^ and F_1_^+^ soils was higher than that of the F_0_^-^ treatment. The FAA–NPR of the F_1_^+^ treatment was the highest (Fig. 7b). Finally, with an increase in nitrogen application, the soil DON content increased (Fig. 7c), and under the same application level, the DON content of the F_0_^-^ treatment was the highest followed by F_1_^+^ and F_0_^+^.

**Fig. 7.**
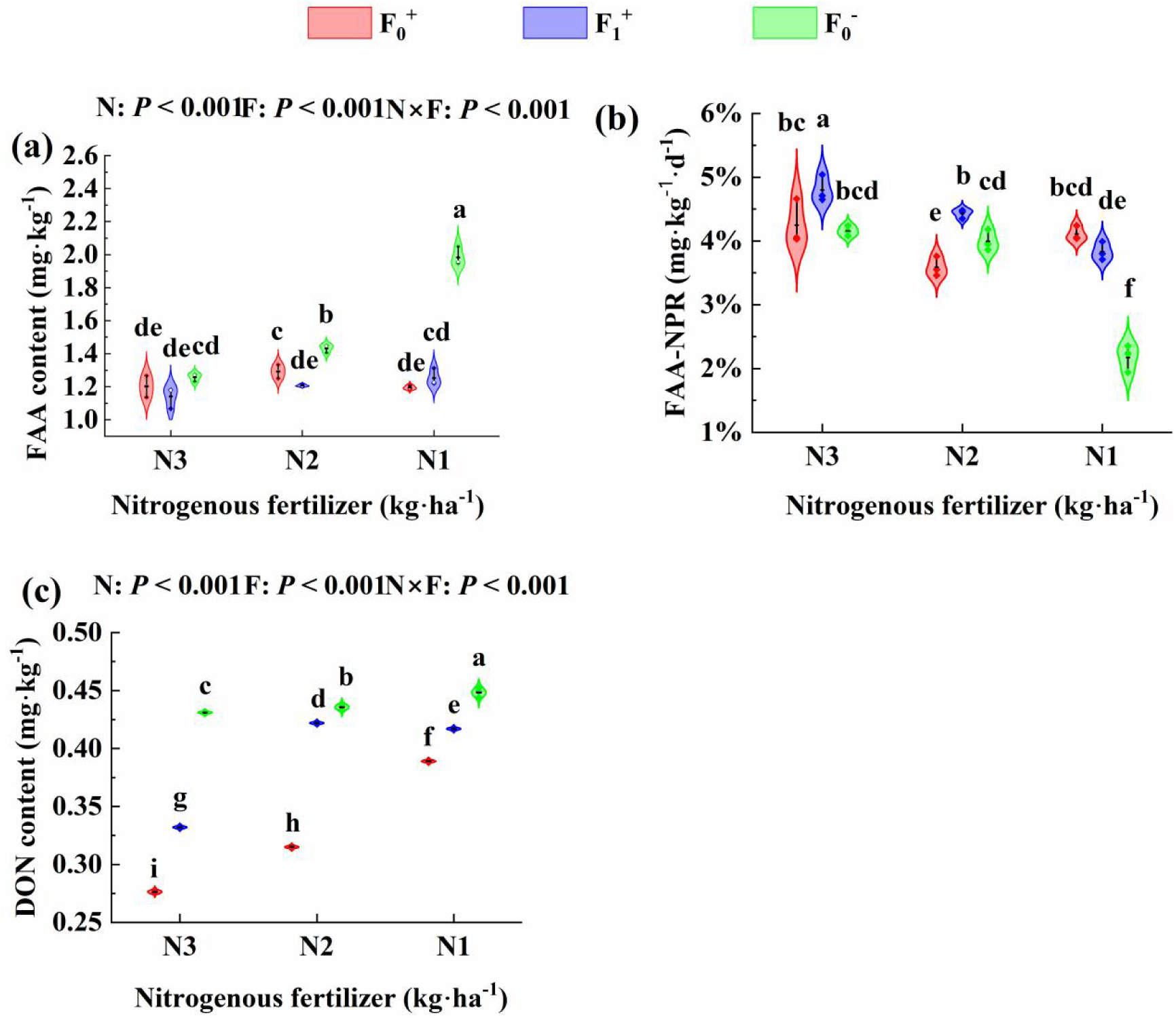
Effects of nitrogen application and AMF on soil (a) free amino acids (FAAs), (b) free amino acid–net production rate (FAA–NPR), and (c) dissolved organic nitrogen (DON).

### Effects of nitrogen application and AMF on soil MBC/MBN, ROC, and IOC

Under the same nitrogen application levels, the soil MBC/MBN treated with F_0_^+^ and F_1_^+^ was higher than that of the soil treated with F_0_^-^. At the N3 level, the MBC/MBN of the F_0_^-^ treatment was the lowest. At the N2 level, there were significant differences among the MBC/MBN for F_0_^+^, F_1_^+^, and F_0_^-^, with the order F_1_^+^ > F_0_^+^ > F_0_^-^. At the N1 level, F_1_^+^ was significantly higher than F_0_^+^ and F_0_^-^ but there was no significant difference between F_0_^+^ and F_0_^-^ (Fig. 8a).

**Fig. 8.**
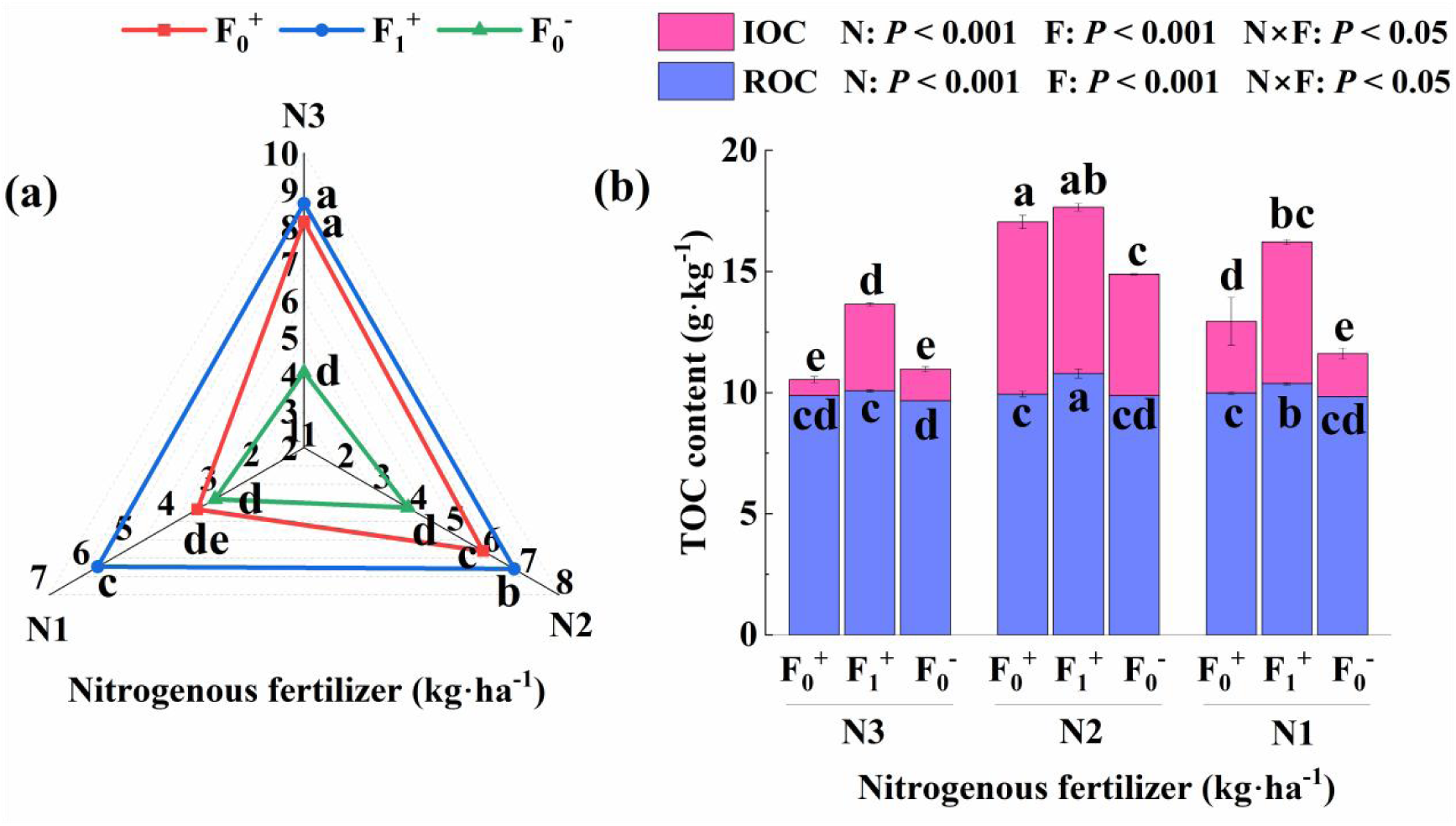
Effects of nitrogen and AMF application on soil MBC/MBN (a) and soil total organic carbon (b). IOC: soil inert organic carbon; ROC: soil reactive organic carbon.

At the N2 level, the total amounts of ROC and IOC in the soil were higher than those at the other nitrogen levels. Among the three nitrogen levels, N2 had the lowest ROC and highest IOC content. Among the three AMF treatments, the F_1_^+^ treatment had the highest IOC and ROC contents, and there was a significant difference between the IOC content of the F_0_^+^ and F_0_^-^ treatments, whereas there was no significant difference in the ROC content between these treatments (Fig. 8b).

### Effects of nitrogen application and AMF application on soil TN and TOC

The soil TN content increased with increasing nitrogen application (Fig. 9a). At the same nitrogen application level, the F_0_^-^ treatment had the highest soil TN content, and the F_1_^+^ treatment had the lowest. Nitrogen application significantly affected the soil TOC content, with the N2 level yielding the highest amount (Fig. 9b). At the same nitrogen level, the relative effect of AMF on soil TOC content was as follows: F_1_^+^ > F_0_^+^ > F_0_^-^.

**Fig. 9.**
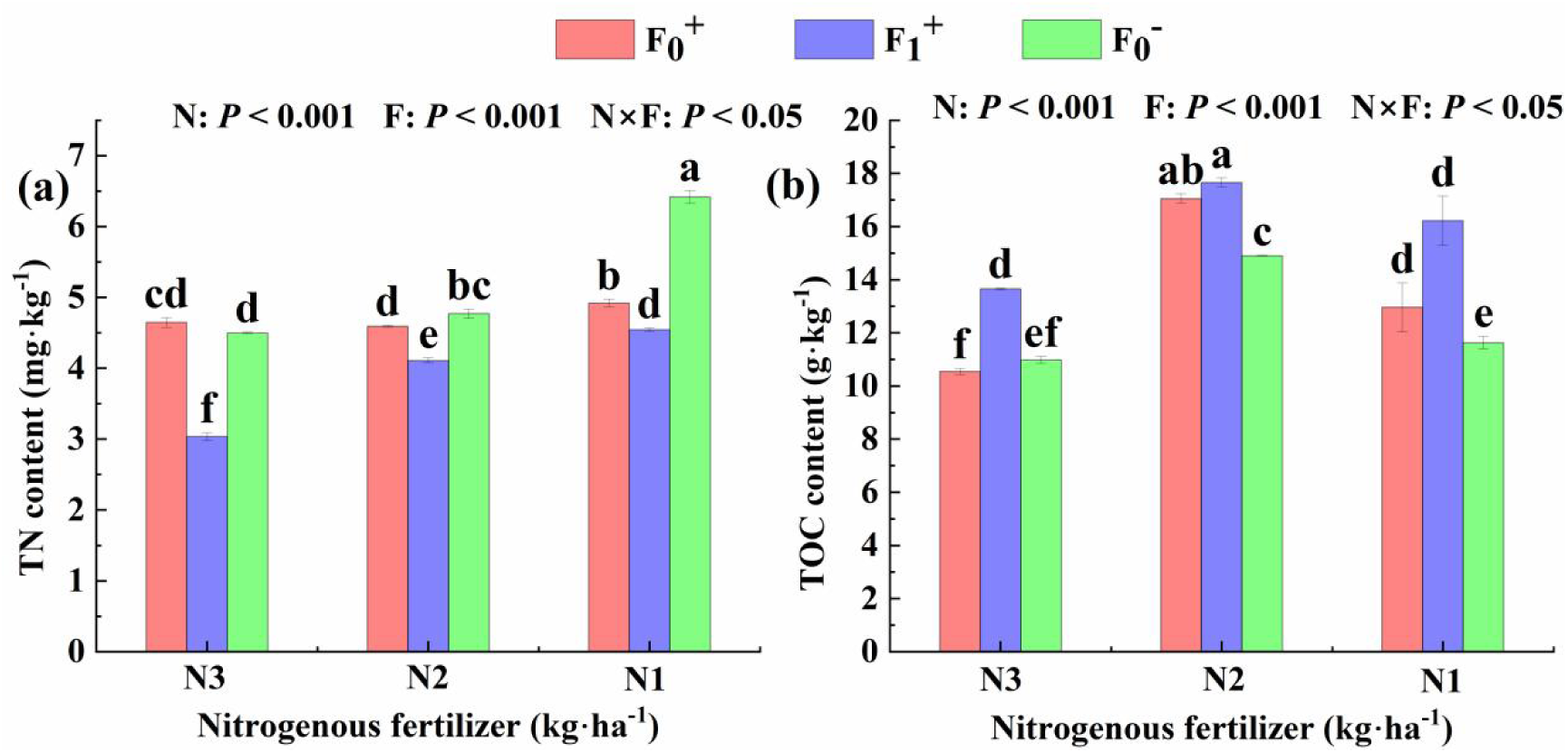
Effects of nitrogen application and AMF on soil (a) total nitrogen (TN) and (b) total organic carbon (TOC). Mean ± standard error.

### Correlations between cotton AMF infection rate, morphological and physiological characteristics, and soil characteristics

There was a significant negative correlation between cotton AMF infection rate and aboveground total nitrogen. Root biomass was negatively correlated with soil TOC but positively correlated with FAA–NPR, plant height, and shoot biomass. Soil protease was positively correlated with N_nit_, whereas PPO was negatively correlated with N_nit_ and N_amo_. There was a significant positive correlation between PPO and the NO_3_^-^-N, NH_4_^+^-N, and DON contents of the soil. Soil NO_3_^-^-N and N_nit_ were also negatively correlated, as were soil NH_4_^+^-N and N_amo_. Soil FAA content was negatively correlated with soil NO_3_^-^-N but positively correlated with N_nit_ (Fig. 10).

**Fig. 10.**
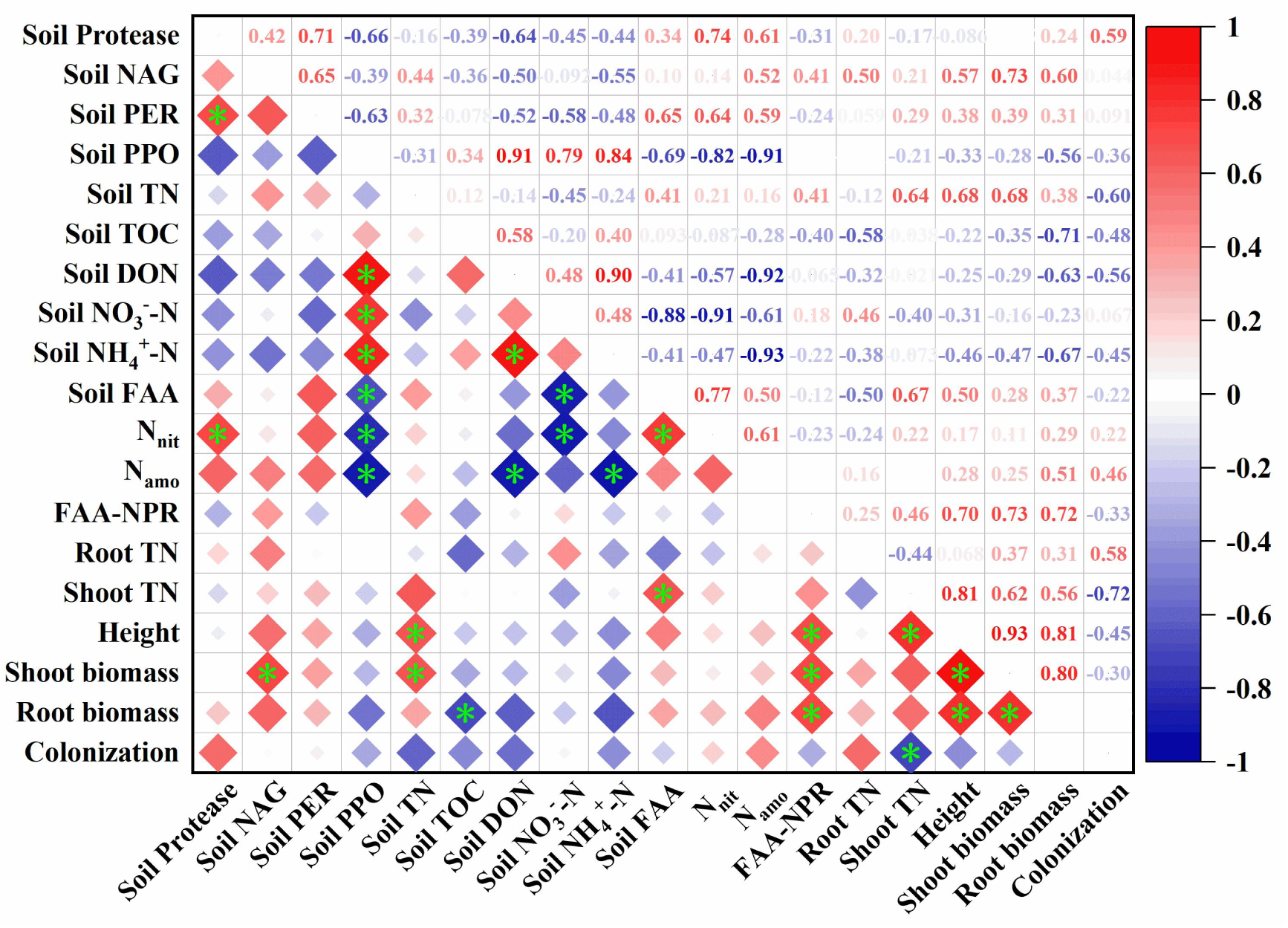
Correlations between cotton AMF infection rate, morphological and physiological characteristics, and soil characteristics (* *P* < 0.05) Note: FAA: Free amino acid; N_nit_: Nitrate nitrogen mineralization rate; N_amo_: Ammonium nitrogen mineralization rate.

## Discussion

Arbuscular mycorrhiza, as the symbiont of soil fungi and plant roots, play an important role in promoting nitrogen uptake by host plants (Lehmann & Rillig, 2015), and the contribution rate of AMF hyphae to plant nitrogen varies between different studies. Using a compartmentalization culture system and isotope-tracer technology, Frey *et al*. (1993) found that the nitrogen absorbed and transported by AMF hyphae to the host plant accounted for 30% of the total nitrogen in the plant. Tanaka and Yano (2005) suggested that up to 75% of the nitrogen in plants is absorbed by AMF hyphae. The results of our study showed that the contribution rate of AMF to plant nitrogen varied with nitrogen application level, at 10.40%, 22.72%, and 16.67% under high, low, and no nitrogen treatments, respectively. This indicates that the application of exogenous nitrogen affects the contribution rate of AMF to cotton nitrogen uptake, and the dependence of cotton on AMF is higher under low-nitrogen conditions. This is consistent with the results of other studies (Zhang *et al*., 2019). Notably, AMF can still contribute nitrogen to cotton under high-nitrogen conditions, but this contribution is relatively low (10.40%). This further indicated that the AMF mycelium was mainly stored by itself, and the input of cotton was secondary under the condition of sufficient exogenous nitrogen. In addition, the AMF infection rate in cotton roots was as high as 83.33% in the absence of N, whereas the contribution rate of AMF to aboveground total N was only 26.66%. Tanaka and Yano (2005) showed that the contribution rate of AMF to aboveground total N in plants could be as high as 74% under such high infection rates. This indicates that the high rate of AMF infection in cotton roots did not result in higher N earnings but instead required cotton to distribute more carbohydrates. Under these conditions, AMF may exhibit “deceptive infection” behavior. Indeed, for the nitrogen reduction level, the soil nitrogen content of the AMF treatment decreased significantly and soil NO_3_^-^-N decreased more than NH_4_^+^-N. This also shows that AMF can promote the absorption and utilization of soil N by cotton, especially NO_3_^-^-N. This results in a decrease in soil nitrogen content.

How can the nitrogen-absorption strategy be adjusted when cotton growth is limited by nitrogen? Root morphology and configuration are adaptive strategies developed by plants for the growth environment (Song & Hou, 2020), which determine the spatial expansion ability of roots and methods of nutrient acquisition (Wang *et al*., 2017). In our study, the colonization of cotton roots by AMF at low-nitrogen levels increased the root surface area, number of tips, and number of branches. This expands the range of root absorption and promotes the ability of the roots to absorb nitrogen directly. In addition, AMF hyphae absorbed more NO_3_^-^-N than NH_4_^+^-N, which is inconsistent with the results of previous studies showing that AMF prefer NH_4_^+^-N (Li *et al*., 2013; Wang *et al*., 2022). This may be related to the N preference of cotton in the study area. The NO_3_^-^-N content of cotton in the study was higher than that of NH_4_^+^-N, and the NH_4_^+^-N content of the soil was higher than that of NO_3_^-^-N. A study conducted by Zhang *et al*. (2023) in this region also supports this conclusion. NO_3_^-^-N is more conducive to root growth and nutrient accumulation, which promotes better root morphology. In addition, the limited uptake of roots at low-nitrogen levels reduces the nutrient concentration in the soil around the roots, leading to the formation of a nitrogen deficiency zone. With the symbiosis between the roots and AMF, the larger biomass and surface area of the hyphae can be extended into the soil beyond the root hair to absorb nitrogen, and the complex hyphal network can accelerate the transport of different forms of nitrogen. This promotes nitrogen nutrition in cotton (Berruti et al., 2016; Run *et al*., 2020). Moreover, the inter-hyphal excitation effect under nitrogen reduction promotes the decomposition of soil nitrogen-containing organic matter and increases the soil organic nitrogen content. This gives cotton plants more options for nitrogen absorption under nitrogen-deficient conditions. Soil NAG, PER, PPO, and proteases are key enzymes for soil nitrogen conversion, of which 4–6% is released into the soil in the form of hyphal secretions (Sinsabaugh, 2010; Sinsabaugh *et al*., 2013). This enhances the activity of soil microorganisms and accelerates the synthesis and release of extracellular soil enzymes. This promotes the conversion of soil nitrogen-containing organic matter into inorganic or simple organic nitrogen (Courty et al., 2010; Jan Jansa *et al*., 2019). In our study, under low-nitrogen conditions, soil NAG, PER, and PPO activities increased significantly, and the rate of nitrogen mineralization also increased. The correlation analysis also showed that soil extracellular enzyme activity was positively correlated with inorganic nitrogen content. These results indicate that AMF promoted the transformation of soil nitrogen, increased the availability of soil nitrogen, and facilitated nitrogen absorption by cotton. According to the traditional nitrogen cycle model, organic nitrogen can be only utilized by plants after it is transformed into inorganic nitrogen by soil free microorganisms, and roots cannot directly obtain organic nitrogen from the soil. In recent years, many studies have shown that plants can directly obtain low-molecular-weight organic nitrogen from soil after symbiosis with AMF, especially soil FAAs (Hobbie & Hobbie, 2008). Studies using isotope-labeling techniques have shown that only AMF mycelia can absorb organic N, which can be quickly transferred to plants (Bukovska *et al*., 2018). In our study, the FAA content in the soil decreased with a reduction in nitrogen application. This indicates that when cotton growth is limited by nitrogen, AMF hyphae may promote the absorption of organic nitrogen, which is then transferred to the root system to meet the nitrogen demand of the cotton plants.

Central to the relationship between roots and AMF is the exchange of carbon (lipids and sugars) and nutrients (Jiang *et al*., 2017). The exchange of carbon and nitrogen between cotton and mycorrhiza is influenced by many factors, especially the soil nutrient status (Johnson *et al*., 1997; Karst *et al*., 2008). When the soil nitrogen supply level is high, the roots can meet the plant needs; therefore, the contribution of the hyphal pathway to nitrogen absorption is small (Chu *et al*., 2020; Zhang *et al*., 2021). When soil nitrogen supply is low, plants preferentially allocate photosynthetic carbon underground to promote nitrogen uptake. Plants are more dependent on the root pathway, hyphal pathway, or both, depending on the tradeoff between carbon investment and nitrogen yield. In our study, the contribution of AMF hyphae to root nitrogen uptake was only 7.92% at low-nitrogen levels. This indicates that nitrogen uptake by cotton is mainly achieved through the root pathway. The reasons for this may be as follows: (1) the current cultivation mode and long-term drip fertilization make cotton develop short and coarse roots, meaning that high investment is required for the plant to attain a high-nitrogen return; (2) in a complex symbiotic environment, the investment in the root system is more stable over time due to the unpredictable environment and unstable resource supply (Zhang *et al*., 2021). In our study, AMF treatment significantly increased the root surface area, number of tips, number of branches, and biomass under the same nitrogen level, and this promoting effect was greater under low-nitrogen conditions. These results indicate that although the contribution rate of AMF mycelia to plant nitrogen was low under the low-nitrogen conditions, AMF may indirectly promote root nitrogen uptake and increase the nitrogen contribution rate by affecting the roots. However, owing to the limitations of the test method, we could not calculate the indirect role of mycelia. Indeed, the contribution of existing mycelia to plant nitrogen was underestimated. AMF infection under the low-nitrogen conditions resulted in coarser roots. According to the cost–benefit theory of root construction (Eissenstat *et al*., 2000), an increase in average root diameter indicates an extension of root life. Moreover, Wu (2022) also showed that moderate nitrogen reduction (272 kg N ha^-1^) can significantly improve root life and productivity. Our study provides evidence explaining the increase in root life under low-nitrogen conditions. Thus, mycorrhizal fungal infection can improve root absorption efficiency and prolong root life under low-nitrogen conditions (Espeleta *et al*., 2009). This may change the tradeoff relationship between root absorption efficiency and root persistence. Although root biomass and AMF infection rates decreased at the higher nitrogen levels, the contribution of AMF to total nitrogen increased. This may be due to the high-nitrogen content in the soil, where the nitrogen storage in the hyphae reaches saturation and surplus nitrogen is transported to the cotton. Secondly, the higher nitrogen level causes the cotton roots to be in a “comfortable state.” The number and elongation range of fine roots were greatly reduced (Zhu *et al*., 2021); however, the death and reconstruction of fine roots take some time, at which point the contribution rate of AMF hyphae to cotton total nitrogen may increase.

Sinsabaugh *et al*. (2013) showed that carbon input into AMF hyphae by plants accounted for 10–20% of plant photosynthetic carbon, of which 6–14% was input into the soil through AMF turnover. The remaining carbon is imported into the mycelium in the form of mycelial secretions, which can be utilized by soil microorganisms, enhancing their activity. In turn, this promotes SOM decomposition (Leigh *et al*., 2011). Our results show that the activity of soil microorganisms was low, even with the effect of AMF, at high-nitrogen levels. This is conducive to the stability of SOM. At low-nitrogen levels, the activity of microorganisms, such as AMF, was enhanced, and the activation effect of the hyphae is conducive to SOM decomposition. This indicates that the exogenous nitrogen application level had a greater effect on soil organic carbon than did AMF. However, previous studies have shown that the activation of hyphae at low-nitrogen levels may promote the formation of stable organic carbon. The potential mechanisms include the following: (1) Low nitrogen accelerates the turnover of microorganisms and increases the amounts of microbial metabolites and residues in the hyphae. (2) Low nitrogen promotes AMF to transfer plant photosynthetic carbon from the rhizosphere hot zone to non-rhizosphere soil, in which microbial activity is low. This results in weak hyphal excitation. (3) Nutrient competition between AMF and soil microorganisms is strong under low-nitrogen conditions, which inhibits the inter-hyphal excitation effect to a certain extent. (4) Low nitrogen stimulates soil microbial activity, improves soil nutrient availability, and promotes host plant growth and rhizosphere deposition. This indirectly increases the input of plant-derived carbon (Brzostek *et al*., 2015; Craig *et al*., 2018). Therefore, the influence of the AMF inter-hyphal excitation effect on soil stable organic carbon depends on the relative extent of the decomposition of the original organic carbon in the inter-hyphal and accumulated microbial residues. This is of great significance for the dynamic cycling of soil carbon and nitrogen.

## Acknowledgements

We express thanks to the Shihezi University Youth Innovative Talents Top program project for supporting this research. We would also like to thank Editage (www.editage.cn) for English language editing.

## Author contributions

W.H.S. developed the idea for research, with extensive discussion with Z.W.F.and P.X.Z. W.H.S. and W.Y.J conducted the glasshouse experiment and analysed its results. C.X.J. and S.Z.H. conducted all analyses relating to soil carbon and infection rate. All authors contributed to the writing and revision of the manuscript.

## Data availability

The datasets used and/or analysed during the current study available from the corresponding author on reasonable request.

